# Trypsin exhibits exopeptidase-like activity toward N-terminal arginine that biases proteomic analyses

**DOI:** 10.64898/2026.05.15.725550

**Authors:** Evan A. Ambrose, Ganapathi Kandasamy, Mia M. Meulenener, Fangliang Zhang

**Author notes:** Lead contact: Fangliang Zhang, Department of Molecular & Cellular Pharmacology; Sylvester Comprehensive Cancer Center, University of Miami Miller School of Medicine, Miami, FL 33136, USA.

## Abstract

Many proteomics protocols rely on enzymatic digestion of complex protein mixtures to generate peptides with predictable cleavage patterns for the mass spectrometry analysis. One of the most utilized enzymes, trypsin, is classically defined as a serine endopeptidase with high specificity for cleaving peptide bonds on the C-terminal side of internal lysine and arginine residues. Accordingly, trypsin is not expected to remove the N-terminal arginine, which may arise through posttranslational modification such as arginylation or by proteolysis exposing internal residues as the new N-termini.

N-terminal arginine plays important biological roles, including functioning as an N-degron and modulating protein interactions/signaling through its positive charge. Curiously, prior mass spectrometry-based studies utilizing trypsin to identify proteins bearing N-terminal arginine have frequently reported low and inconsistent yields, suggesting potential systematic bias in current proteomic approaches.

Here, we explored whether trypsin would affect the integrity of the N-terminal arginine. By using antibodies specifically recognizing N-terminal arginine of different peptides, and by using mass spectrometry peptide analysis, we show that trypsin can remove N-terminal arginine residues in an exopeptidase-like manner. This effect occurs across a range of digestion conditions consistent with standard proteomic workflows, on peptides or whole proteins, and depends on trypsin concentration, incubation time, and catalytic activity. In addition, we show that the alternative arginine-cleavage enzyme Arg-C can also affect N-terminal arginine in a sequence-dependent context. In contrast, Lys-C and LysargiNase do not exhibit such effects, providing suitable alternative digestion strategies.

Together, these findings reveal an unappreciated enzymatic behavior of arginine-cleaving proteases and suggest that their widespread use may systematically compromise the detection of N-terminal arginine in proteomic studies.

## INTRODUCTION

Large-scale proteomic studies typically utilize a bottom-up approach, in which complex protein mixtures are enzymatically digested into constituent peptides before detection and identification by mass spectrometry (MS)^1,2^. Among available proteases, trypsin is the predominant choice. Trypsin, as classified by the International Union of Biochemistry and Molecular Biology, is a serine endopeptidase which specifically cleaves peptide bonds on the C-terminal side of arginine and lysine residues^3–5^. The use of trypsin generates peptides with favorable physicochemical properties, including predictable fragmentation patterns, efficient electrospray ionization, and an optimal mass range for tandem MS (MS/MS). However, the near-ubiquitous use of trypsin may also introduce analytical bias, particularly when the biological question of interest could be masked by the activity of trypsin. One such consideration is the analysis of proteins or peptides containing N-terminal arginine, for which there is an emerging concern that it could be affected by trypsin.

The N-terminal arginine often functions as part of a type I N-degron, signaling for rapid ubiquitin-dependent turnover^6–9^. In other contexts, N-terminal arginine contributes to regulatory functions independent of degradation^10–12^. N-terminal arginine-containing proteins can arise from multiple sources. A major source is a post-translational modification termed arginylation^13–15^. This reaction is catalyzed by the highly conserved enzyme arginyltransferase (Ate) in the majority of eukaryotes, and by Ate-like proteins (AteL) in the other eukaryotic clades^16–18^. Another source is from the actions of proteases. For example, certain types of calpains favor arginine at position P1’ in the recognition site, leading to exposure of arginine on the N-terminus of the protein fragment after cleavage^19–21^. Caspases can also cleave proteins to expose N-terminal arginine, although at a relatively low frequency^22–24^. The identification of proteins or protein fragments containing N-terminal arginine bears high biological significance. Arginylation is found in nearly all eukaryotes. The dysregulation of arginylation or Ate has been implicated in various disease states including cancer^25–29^, cardiovascular disorders^11,30–33^, metabolic stress responses^34–38^, tissue injury^39–42^, and development abnormalities^43–46^. Consistently, evidence suggests that the N-terminal arginine on proteolytic fragments serves important roles in cell death regulation and other cellular functions^20,47–49^. For these reasons, systematic characterization of protein or protein fragments containing N-terminal arginine is essential for understanding this regulatory system.

Biochemical studies with engineered peptides have suggested the presence of a significant number of proteins or protein fragments with N-terminal arginine *in vivo*, either by arginylation or by proteolysis. In the case of Ate-catalyzed arginylation, it was predicted that as high as 25% of the proteome may be subjected to this modification^50,51^. Despite this broad potential, comprehensive identification of arginylated proteins in complex biological samples has proven challenging. Several MS-based approaches have attempted higher-throughput discovery, yet these studies have produced relatively small and poorly overlapping sets of hits, suggesting substantial technical limitations in detecting N-terminal arginine by standard proteomic workflows.

Multiple experimental efforts illustrate this problem. Currently, all the demonstrated attempts using MS to identify proteins with N-terminal arginylation had to use either preferential labeling and/or enrichment. Among notable examples, Decca, et al., labeled proteins with ^14^C arginine in a translation-free system, separated using 2D gel electrophoresis, which resulted in the identification of 6 proteins by matrix-assisted laser desorption/ionization time-of-flight (MALDI-TOF)^52^. In another study by Wong and Xu, et al., antibody-based enrichment targeting N-terminal RD- or RE-containing proteins expanded to the detectable set to 43 proteins by liquid chromatography coupled tandem mass spectrometry (LC-MS/MS)^53^. More recently, ex vivo assays using ribosome-depletion and Ate over-supplementation were employed to maximize the separation of posttranslational arginylation and translational incorporation of arginine^54^. This approach yielded 235 arginylation sites, a number that is higher than previous attempts but remains far below the scope predicted from biochemical principles^50,51^. In addition, the sets of arginylated proteins identified in different studies exhibit little overlap. Even when the same proteins are detected across studies, the indicated sites of arginylation frequently differ. Moreover, several well-established arginylation targets identified through low-throughput biochemical techniques, such as RGS proteins initially identified by Edman degradation^55,56^, consistently failed to appear in MS-based screenings. The lack of robustness was further exemplified in a recent study by Seo, et al, in which the researchers utilized the ZZ domain of p62 to pull-down proteins containing N-terminal arginine^57^. While this study identified 59 proteins in the pulldown fraction, their identities were solely determined through the sequences of internal peptides and not by any direct demonstration of the existence of the N-terminal arginine on these proteins by MS.

This persistent lack of sensitivity and robustness of MS-based method for detecting N-terminal arginine of proteins is in stark contrast to numerous MS-based studies that have positively identified proteins containing other types of N-terminal residues in vivo^58–62^. Part of this problem may be attributed to the unstable nature of proteins with N-terminal arginine, which could reduce the abundance of such protein species even in the presence of proteasome and autophagy inhibitors due to their leaky nature^7,9,63,64^. However, the root of this problem may run deeper, particularly considering the persistence of the same issue even when arginylated species are pre-enriched through antibodies or specific binding domains^53,57^. This discrepancy raises the possibility that the sample-processing steps used in standard bottom-up proteomics may themselves contribute to the loss of N-terminal arginine.

Trypsin is traditionally regarded as a strict endopeptidase that does not remove N-terminal residues, making it the default enzyme in most high-throughput proteomic workflows, including all the previous efforts in the investigation of arginylation^50–54,57,65^. However, the consistent absence of N-terminal arginine-containing peptides in MS led us to hypothesize that trypsin may possess a previously overlooked exopeptidase-like activity capable of removing N-terminal arginine under certain conditions.

In the present study, we systematically tested the potential impacts of trypsin treatment on N-terminal arginine residue. By using multiple arginylation-specific antibodies, MALDI-MS, and MS-MS, we demonstrate that trypsin digestion results in a concentration- and time-dependent loss of N-terminal arginine, with a dependency on the enzymatic activity of trypsin. Additionally, we find that Arg-C, another commonly used protease in proteomics workflows, exhibits a similar risk of N-terminal arginine loss, while two other proteases (Lys-C and LysargiNase) may constitute suitable alternatives. Together, these results reveal an unrecognized limitation in conventional MS-based approaches for identifying arginylated proteins. They underscore the need to reevaluate prior datasets and to adopt alternative digestion strategies when studying N-terminal modifications such as arginylation. Furthermore, this discovery may indicate incomplete characterization of N-terminal arginine (or lysine) arising from other cellular processes, such as cleavage by proteases. By highlighting this enzymatic vulnerability inherent to trypsin, our work provides a mechanistic explanation for the longstanding difficulty in identifying N-terminally arginylated proteins and offers a path forward for improving proteomic detection of this important post-translational modification.

## RESULTS

### Trypsin removes the signal of N-terminal arginine of peptides recognized by specific antibodies

We previously generated multiple antibodies specific to peptides containing N-terminal arginine (Figure 1A). These include an antibody against RAA15 (RAAVAAENEEIGAHIKHC, derived from arginylated Talin-1 fragment)^12^, R-HIF (REGAGGENEKKKMSC, derived from arginylated hypoxia inducible factor 1α (Hif1α))^18^, RDD11 (RDDIAALVVDC, derived from arginylated beta actin)^10^, and REHQ (REHQLLC). These antibodies were validated for their specificities to detect corresponding peptides containing N-terminal arginine, while exhibiting minimal cross-reactivity to the respective non-arginylated peptides.

**Figure 1.**
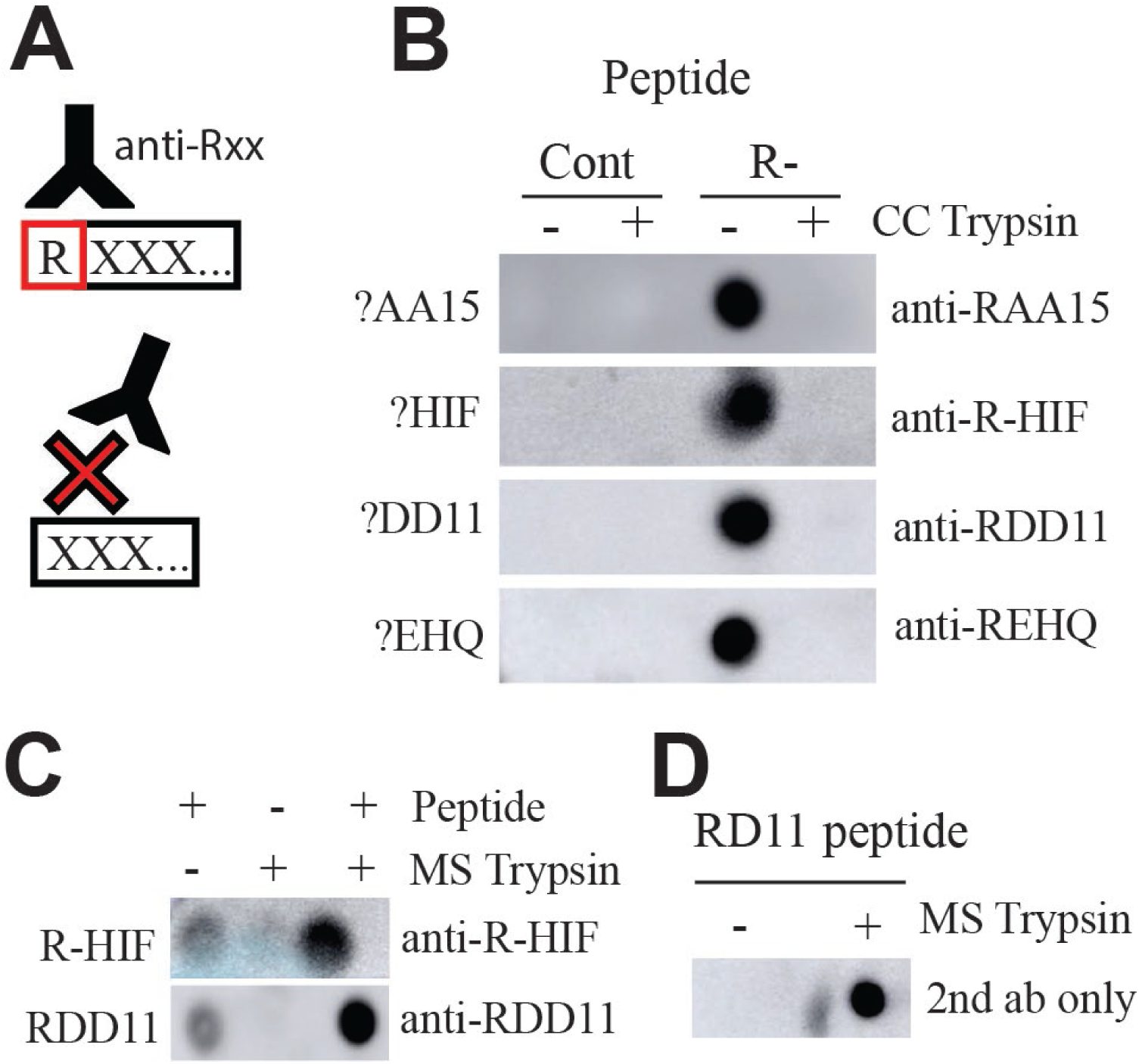
Trypsin removes signal of N-terminal arginine recognized by specific antibodies. A) Antibodies were generated to specifically recognize N-terminally arginylated peptides including RAA15, R-HIF, and RDD15, which were derived from the sequences of the arginylated Talin-1 fragment, Hif1α, and beta actin, respectively. These antibodies were validated to have minimum cross-reactivity with the non-arginylated peptide controls. B) The 4 different N-terminally arginylated peptides and their corresponding non-arginylated controls, together termed as ?AA15 (RAA15 or AA15), ?HIF (R-HIF or HIF), ?DD11 (RDD11 or DD11), and ?EHQ (EHQ or REHQ) at final concentrations of 75 ng/μL, were incubated with cell-culture grade (CC) trypsin (2.1 ug/μL) at 37°C for 18 hours. The reaction was terminated by denaturing at 100°C for 30 minutes. Then, 1.5μL of the reaction mix was probed by dot blotting with the corresponding arginylation-specific antibodies (anti-RAA15, anti-R-HIF, anti-RDD11, and ant-REHQ antibody)). C) Similar to B, except that the two arginylated peptides were incubated with mass spectrometry-grade (MS) trypsin (1 μg/μL). Increased signals were observed with trypsin present compared to the trypsin-absent controls, indicating the MS-grade trypsin is not compatible with the dot blotting assay. The exposure was reduced to illustrate the difference in signal intensity between untreated and MS trypsin-treated samples. D) Arginylated RDD11 (75 ng/μL) was incubated with MS-grade trypsin (1 μg/μL) with conditions similar to B, except that the dot blot was probed with the secondary antibody alone (anti-Rabbit-HRP) without the primary antibody.

To gain a first impression on whether trypsin can remove N-terminal arginine from peptides, we utilized the RAA15, R-HIF, RDD11, and REHQ peptides together with the matching control non-arginylated peptides. These peptides were incubated with trypsin (0.21%, cell-culture grade for the initial tests) at 37°C overnight. Note that in this condition the molar ratios of trypsin to peptide substrates were ∼1:1-2 and within the upper limit of usable MS digestion^66–68^. The resulting product was heat inactivated and then probed with corresponding arginylation-specific antibodies by dot blotting (Figure 1B). We found that the N-terminal arginylated peptide signal disappeared with trypsin treatment for all four pairs of peptides tested, potentially indicating the arginine had been removed. We attempted to validate these results with MS-grade trypsin, which is treated with L-(tosylamido-2-phenyl) ethyl chloromethyl ketone to inactivate chymotrypsin activity and has lysine side chains acetylated to reduce autolysis. However, the presence of MS-grade trypsin led to increased antibody signal compared to peptides in buffer alone (Figure 1C). In some instances, the application of the MS-grade trypsin was sufficient to produce a strong signal with the secondary antibody alone (Figure 1D). These results indicate that MS-grade trypsin has non-specific reactivity with the antibodies and is therefore not suitable for immunoblotting. Thus, all subsequent antibody-based experiments employed cell-culture grade trypsin (CC trypsin).

To determine whether the enzymatic activity of trypsin is required for the arginine removal, trypsin was heat inactivated prior to incubation with the peptide substrates. We found that pre-digestion heat inactivation preserved N-terminal arginine signal (Figure 2A). In comparison, a post-incubation heat treatment did not generate such an outcome (Figure 2B). These data together suggest that the loss of the N-terminal arginine during trypsin treatment is not due to spontaneous hydrolysis. To further determine whether this effect depends on the enzymatic activity of trypsin, trypsin was inhibited with trypsin neutralizing solution containing competitive trypsin inhibitor prior to incubation with peptide substrates. In this case, the loss of N-terminal arginine signal was attenuated (Figure 2C). These results indicate that the removal of N-terminal arginine requires the catalytic activity of trypsin, supporting an enzymatic basis for this exopeptidase-like behavior rather than nonspecific degradation.

**Figure 2.**
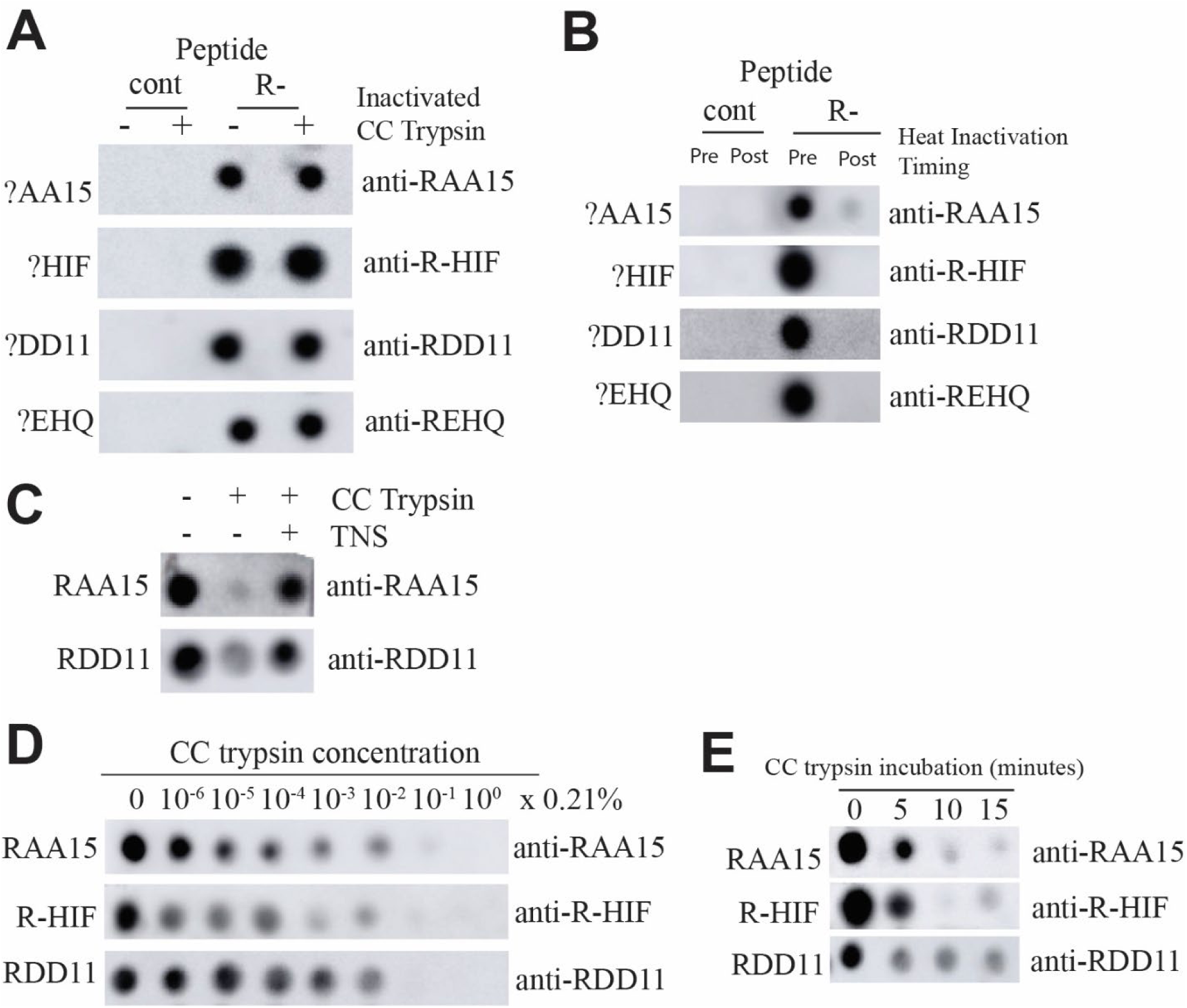
Characterization of trypsin exopeptidase activity. A) The 4 different arginylated peptides (“R-”), RAA15, R-HIF, RDD11, or REHQ (75 ng/μL), and the corresponding non-arginylated control (“cont”), together referring as “?AA15”, “?HIF”, “?DD11”, and “?EHQ””, were incubated with heat-inactivated cell-culture grade (CC) trypsin (2.1 μg/μL) or buffer alone and probed by dot blotting with 1.5μL of the reaction and detection with the corresponding arginylation-specific antibodies. B) Similar to A, except that the heat inactivation (100°C for 30min) was performed either in the beginning (“Pre”) or the end (“Post”) of the incubation with CC trypsin (2.1 μg/μL). C) Arginylated peptides RAA15 or RDD11 (75 ng/μL) were incubated for 30 minutes with CC trypsin (0.04% final concentration) or HBSS buffer. Trypsin was pre-treated for 15 minutes with trypsin neutralizing solution (TNS)at ratio of 1:4 or 1:1(trypsin vs. TNS) for RAA15 and RDD11, respectively, or with equivalent volume of HBSS buffer. The resulting products were probed by dot blotting and detected with the corresponding arginylation-specific antibodies. D) 3 arginylated peptides were incubated with a series of 10-fold dilutions of CC trypsin (starting at 2.1 μg/μL) and probed by arginylation-specific antibodies. E) 3 arginylated peptides were incubated with CC trypsin (0.21 μg/μL). The reaction was stopped at the indicated times by heat inactivation and probed by dot blotting.

To characterize the kinetics of trypsin digestion, a dose-response curve was generated using ten-fold dilutions of trypsin with either RAA15, R-HIF, or RDD11 peptide at 37°C overnight followed by heat inactivation (Figure 2D). While a complete disappearance of the N-terminal arginine signal was observed at 0.021% trypsin, a non-negligible reduction can be observed at lower concentrations such as 0.00021%, which is ∼10x times below typical trypsin concentrations used in most MS applications (∼0.002% trypsin assuming 1mg/mL of protein and 1:50 w/w)^69^. Additionally, the N-terminal arginine signals of different peptides appear to have different sensitivities towards trypsin, potentially indicating sequence-dependent variance. To determine the time-dynamics, the reaction using 0.021% trypsin was stopped at intervals ranging from 0 to 15 minutes for the different peptides (Figure 2E). Nearly all the arginine-containing peptide signal was depleted within 15 minutes. These results indicate that trypsin is highly effective in removing N-terminal arginine.

Together, these results indicate that trypsin likely can remove N-terminal arginine from peptides, as was observed with multiple peptides. However, these do not remove the possibility of nonspecific cleavage, and more direct evidence is required to conclude cleavage is occurring at the N-terminal arginine.

### Mass spectrometry confirms the removal of N-terminal arginine by different grades of trypsin

To validate results on immunoblots, and to ensure that the results with cell-culture grade trypsin were reproducible with MS-grade trypsin, N-terminally arginylated RAA15 and R-HIF peptides were treated with either grade of trypsin and analyzed by MALDI-TOF in MS (Figure 3, 4). When incubated with trypsin, the expected mass of the peptide without arginine was identified, along with peaks indicating cleavage at the internal lysine residue. Results were consistent with both grades of trypsin. Notably, the cleavage was biased towards the internal lysine instead of the N-terminal arginine, although the ratio differed among peptides, suggesting the efficiency of N-terminal cleavage may be affected by peptide sequence.

**Figure 3.**
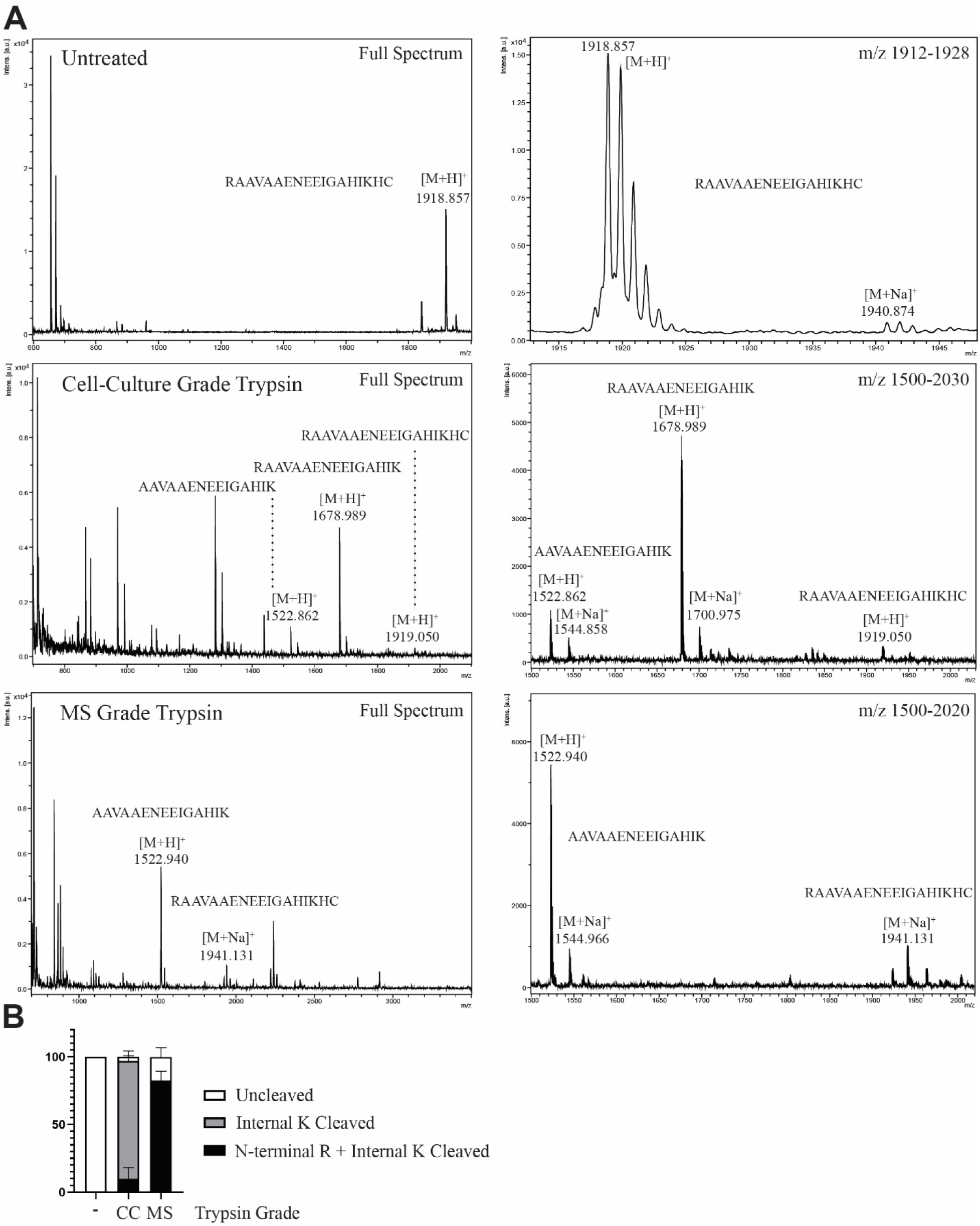
MADLI-TOF demonstrates removal of N-terminal arginine from RAA15 peptide by different grades of trypsin. A) The RAA15 peptide (75 ng/μL) was incubated with cell-culture grade (2.1 μg/μL) or mass-spectrometry (MS) grade trypsin (1 μg/μL) for 18 hours. The reactions were terminated by heat inactivation and then analyzed by MALDI-ToF for the composition of the peptides. B) Quantification of the ratios of ion peaks representing arginylated or non-arginylated forms of the peptides in different treatment conditions. The measurements were performed in two biological repeats with technical triplicates in each run. Error bars indicate standard deviation.

**Figure 4.**
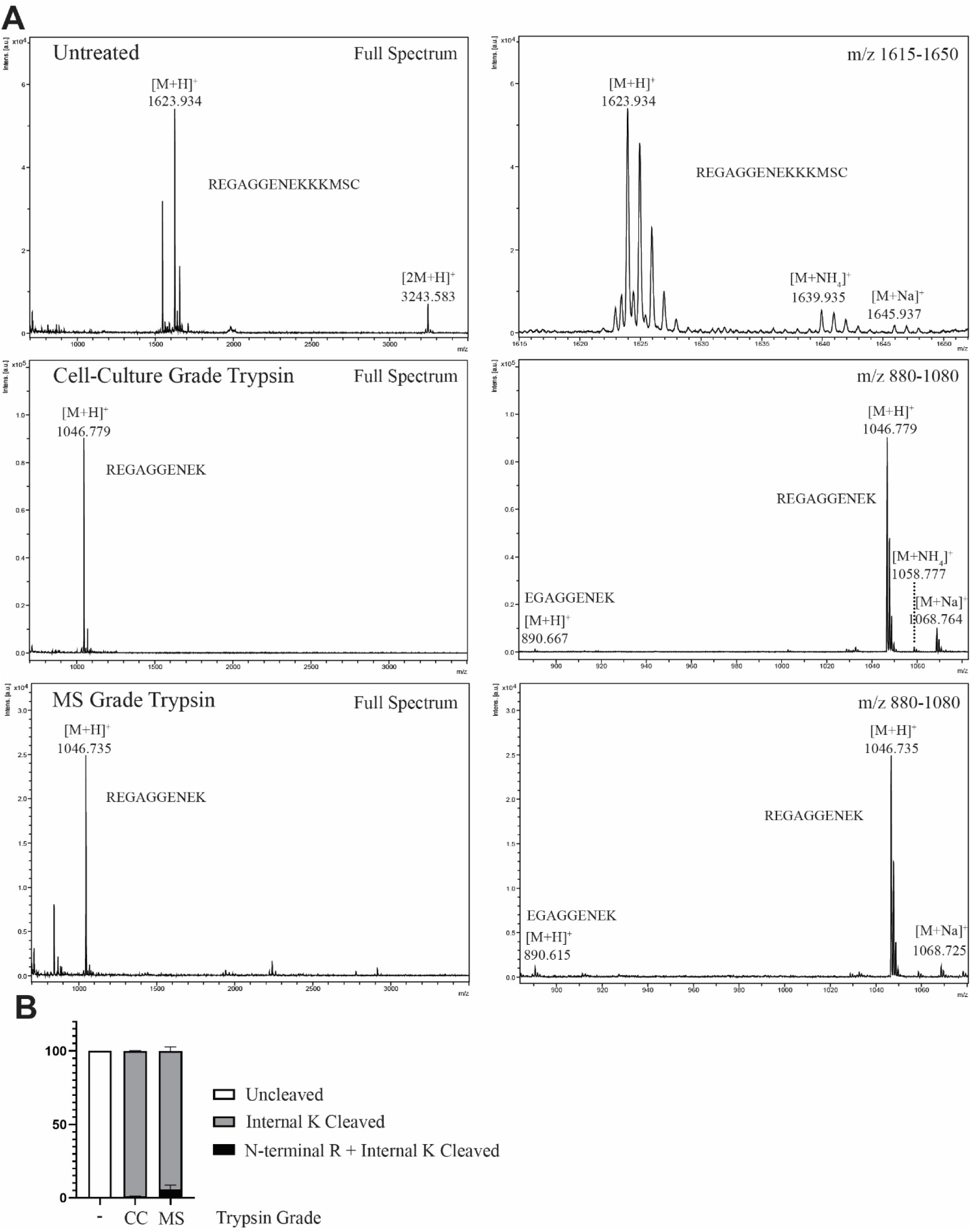
MADLI-TOF demonstrates removal of N-terminal arginine from R-HIF peptide by different grades of trypsin. A) The R-HIF peptide (75 ng/μL) was incubated with cell-culture grade (2.1 μg/μL) or mass-spectrometry (MS) grade trypsin (1 μg/μL) and analyzed by MALDI-ToF. The reactions were terminated by heat inactivation and then analyzed by MALDI-ToF for the composition of the peptides. B) Quantification of the ratios of ion peaks representing arginylated or non-arginylated forms of the peptides in different treatment conditions. The measurements were performed in two biological repeats with technical triplicates in each run. Error bars indicate standard deviation.

To obtain further confirmation of the removal of the N-terminal arginine, LC-MS/MS analysis was conducted on one of the peptides (R-HIF) treated with cell-culture grade (Figure 5B,C) or MS-grade trypsin (Figure 6). The test revealed the presence of peptide with the N-terminal arginine removed when incubated with trypsin, which was not observed in the absence of trypsin. MS/MS confirmed the sequence of R-HIF with N-terminal arginine removed. Therefore, both MS techniques confirm the ability of trypsin, either cell culture grade or MS grade, to cleave N-terminal arginine from peptides.

**Figure 5.**
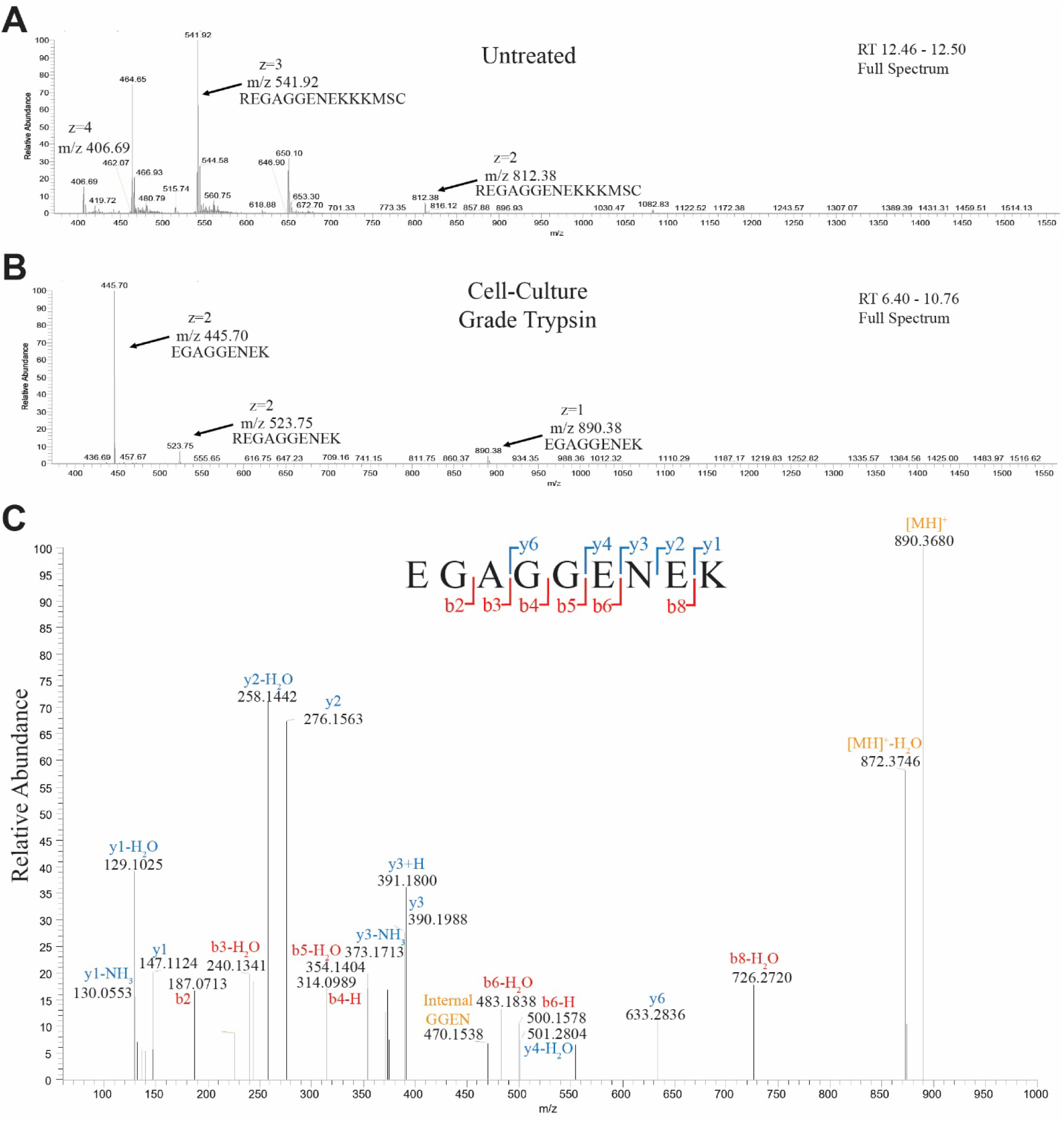
LC-MS/MS confirms N-terminal arginine cleavage by cell-culture grade trypsin. A,B) MS1 spectra of samples restricted to the indicated retention time range from UHPLC input. The R-HIF peptide (75 ng/μL) was incubated with buffer (A) or cell-culture grade trypsin (2.1 μg/μL) (B) and analyzed by LC-MS/MS. The peaks representing the non-cleaved form, the form with internal cleavage, and the form with N-terminal cleavage were indicated. C) Representative MS/MS spectrum showing the expected ion peaks from a sequence of R-HIF peptide with N-terminal arginine removed. Some fragment ions differed by ±1 Da from predicted neutral masses, consistent with expected protonation/deprotonation states of b-and y-ions during fragmentation.

**Figure 6.**
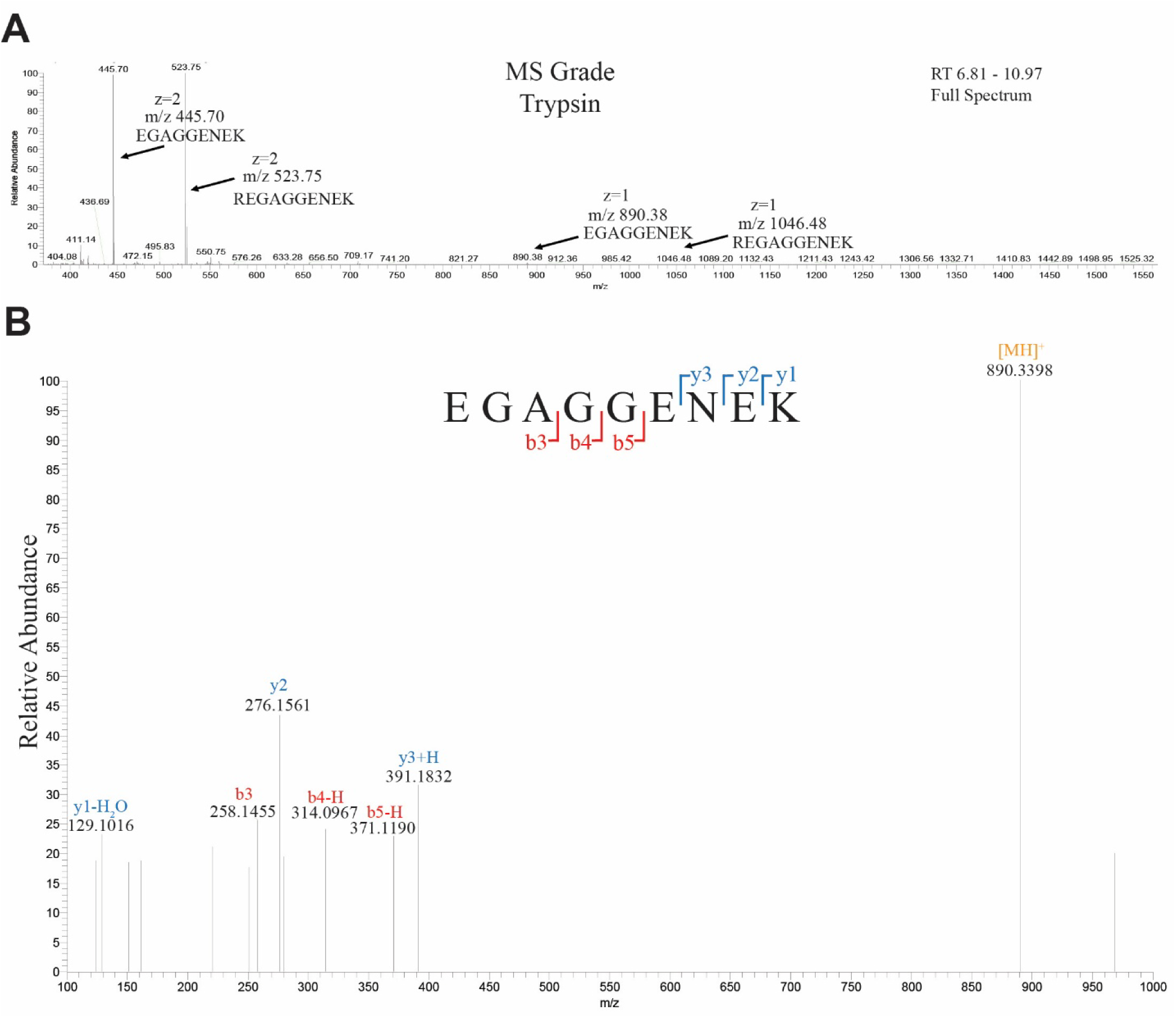
LC-MS/MS confirms N-terminal arginine cleavage by mass-spectrometry grade trypsin. A) MS1 spectrum of sample restricted to the indicated retention time range from UHPLC input. R-HIF peptide (75 ng/μL) was incubated mass-spectrometry grade trypsin (1 μg/μL) and analyzed by LC-MS/MS. The peaks representing the form with internal cleavage and the form with N-terminal cleavage were indicated. B) Representative MS/MS spectrum of R-HIF peptide with N-terminal arginine removed. Some fragment ions differed by ±1 Da from predicted neutral masses, consistent with expected protonation/deprotonation states of b- and y-ions during fragmentation.

### Trypsin cleaves N-terminal arginine from entire proteins and with other proteins present

Samples for proteomic experiments often contain thousands of proteins, instead of a singular peptide. To determine whether trypsin could cleave N-terminal arginine in a solution containing multiple proteins, casein (4.25mg/mL final concentration) was added to the reaction of either RAA15 or R-HIF peptides and cell-culture grade trypsin (∼1ug/uL) (Figure 7A). Dot blots showed a nearly complete loss in arginylation signal using specific antibodies, indicating N-terminal arginine was removed even in the presence of other proteins. To ensure trypsin was active in these reactions, remaining samples were analyzed by SDS-PAGE and Coomassie staining, showing cleavage of casein treated with trypsin. These results suggest that trypsin is able to remove N-terminal arginine in protein mixture.

**Figure 7.**
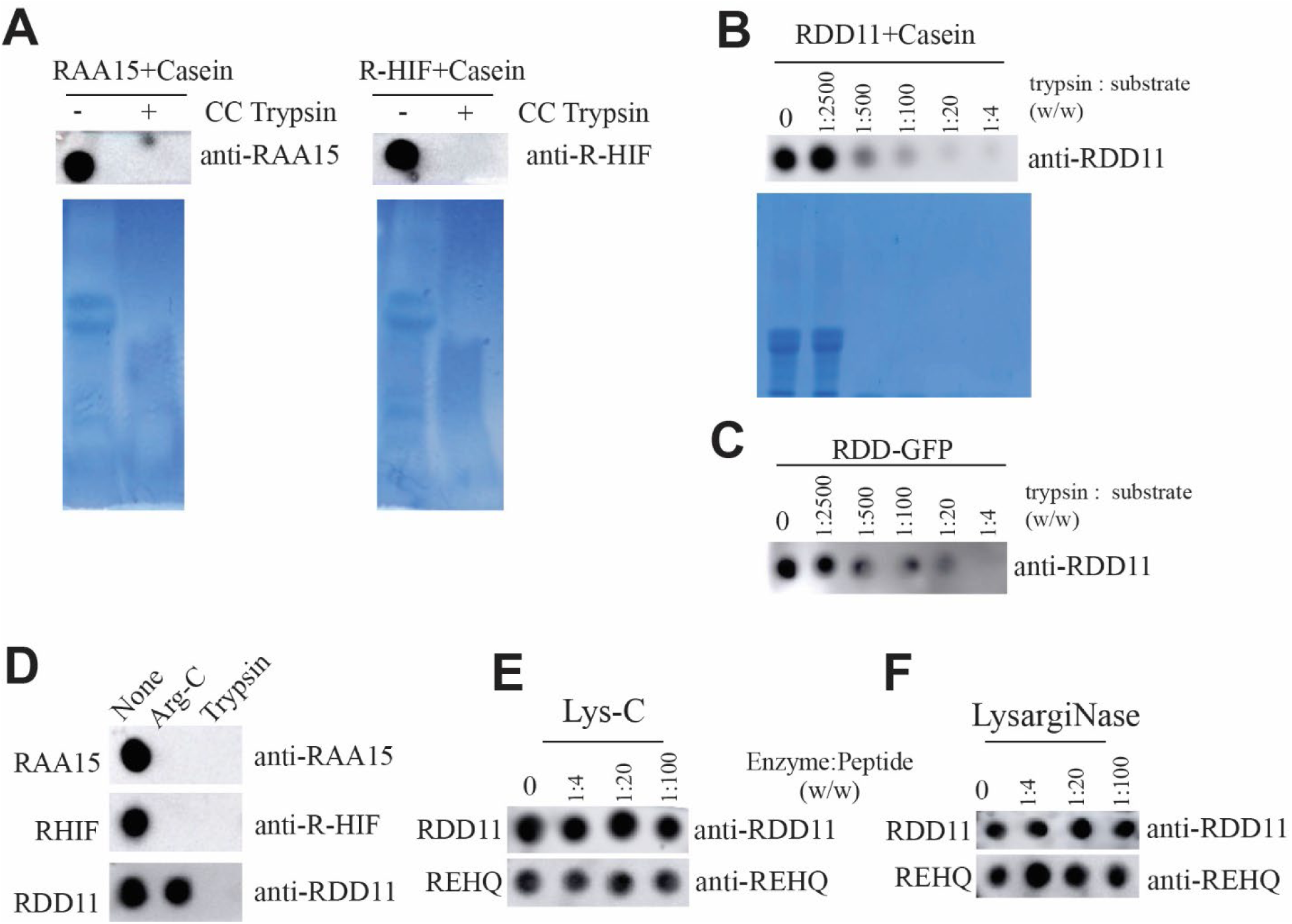
Trypsin cleaves N-terminal arginine in conditions similar to typical proteomics experiments. A) RAA15 or R-HIF peptide (75 ng/μL final concentration) mixed with casein (4.25 μg/μL final concentration) was incubated with cell-culture grade trypsin (1μg/μL final concentration) at 37°C for 18 hours before denaturation at 100°C for 30 minutes. The reaction products were probed by dot blotting with corresponding arginylation-specific antibodies. The same products were analyzed by SDS-PAGE for cleavage of casein proteins. B) RDD11 peptide (75 ng/μL final concentration) mixed with casein (4.25 μg/μL final concentration) was incubated with 5-fold dilutions of cell-culture grade trypsin from 1:2500 to 1:4 (w/w) at 37°C for 18 hours before denaturation at 100°C for 30 minutes. The reaction products were probed by dot blotting with the anti-RDD11 antibody. The same products were analyzed by SDS-PAGE for cleavage of casein proteins. C) Purified RDD-GFP recombinant protein (175 ng/μL) was incubated with trypsin of a series of 10-fold dilutions starting at 2.1 μg/μL. The resulting products were probed by dot blotting with an arginylation-specific antibody. D) 3 different arginylated peptides RAA15, R-HIF, and RDD11 (75 ng/μL) were incubated with ArgC (425 ng/μL) or trypsin (2.1 μg/μL) and probed by dot blotting with corresponding arginylation-specific antibodies. E) Two different arginylated peptides, RDD11 and REHQ (75 ng/μL) were incubated with 5-fold dilutions of LysC from 1:4 to 1:100 (w/w) at 37°C for 18 hours before denaturation at 100°C for 30 minutes. The reaction products were probed by dot blotting with anti-RDD11 or anti-REHQ antibodies. F) Two different arginylated peptides, RDD11 and REHQ (75 ng/μL) were incubated with 5-fold dilutions of LysargiNase from 1:4 to 1:100 (w/w) at 37°C for 18 hours before denaturation at 100°C for 30 minutes. The reaction products were probed by dot blotting with anti-RDD11 or anti-REHQ antibodies.

To more closely approximate real-life proteomic conditions with mixed protein samples, RDD11 peptide was mixed with casein and incubated with a series of trypsin concentrations spanning 1:2500 to 1:4 (trypsin:substrate, w/w) (Figure 7B). We found that the N-terminal arginine signal was progressively reduced across this range, with substantial loss observed within concentrations commonly used in proteomic workflows (starting at 1:500). To further determine whether trypsin could remove N-terminal arginine from an entire protein, we used RDD-GFP, a recombinant protein containing a N-terminal sequence identical to the RDD11 sequence fused to a GFP. A noticeable reduction in N-terminal arginine signal was observed at ∼1:500, with near-complete loss occurring at ∼1:20 (Figure 7C).

These results indicate that trypsin-mediated removal of N-terminal arginine occurs under physiologically relevant digestion conditions and is not restricted to high enzyme concentrations.

### Alternative digestion enzymes for the analysis of N-terminal arginine

To identify proteases compatible with the detection of N-terminal arginine, we evaluated additional enzymes commonly used to cleave arginine and lysine residues in mass spectrometry workflows. These include Clostripain (Arg-C), Endoproteinase Lys-C (Lys-C), and LysargiNase. Arg-C and Lys-C specifically cleave proteins on the C-terminus of arginine or lysine residues, respectively, while LysargiNase cleaves on the N-terminus of arginine and lysine. Thus Arg-C together with Lys-C, or LysargiNase alone, can be used for MS analysis as an alternative to trypsin^70,71^.

Similar to trypsin, exopeptidase activity has not been reported for ArgC. To test this possibility, Arg-C (425ng/uL) was incubated with either RAA15, R-HIF, and RDD11 peptides overnight and the presence of N-terminal arginine was probed by dot blotting with corresponding antibodies (Figure 7D). We found that the presence of Arg-C led to complete disappearance of N-terminal arginine signal for RAA15 and R-HIF peptides, indicating that the arginine was likely cleaved. However, N-terminal arginine signal was unaffected by Arg-C when a different peptide, RDD11, was used. These results indicate ArgC may exhibit sequence-specific removal of N-terminal arginine and is likely not appropriate in high-throughput screening. In contrast to trypsin and Arg-C, neither Lys-C nor LysargiNase led to a detectable loss of N-terminal arginine signal over different peptides, indicating that these enzymes do not exhibit exopeptidase-like removal (Figure 7E&F). These findings suggest that Lys-C and LysargiNase may represent suitable alternatives for proteomic analyses when preservation of N-terminal arginine is required.

## Discussion

Trypsin has long been regarded as a classic digestive serine protease with highly defined endoproteolytic specificity. This predictable behavior has made trypsin the gold-standard protease in proteomics, where its reliability and well-understood specificity are foundational assumptions in experimental design and data interpretation. In this context, trypsin has generally been treated as mechanistically constrained, with its biochemical properties considered fully understood. Our findings challenge this long-standing view and reveal that the functional repertoire of trypsin is broader than previously appreciated.

Specifically, trypsin was not considered a risk factor for the integrity of N-terminal arginine residues, which can arise from post-translational arginylation or from proteolytic processing by endogenous proteases. Consequently, trypsin has been widely used for the identification of proteins bearing N-terminal arginine in past investigations. Here, our study indicates that trypsin exhibits a previously unrecognized exopeptidase-like activity that can remove N-terminal arginine residues. While we do not formally establish the enzymatic mechanism or kinetics underlying this activity, the observed removal of N-terminal arginine across multiple independent assays overturns a core assumption in standard proteomics workflows, suggesting that trypsin may not be suitable for analyses when preservation or accurate identification of N-terminal arginine is required.

The exopeptidase-like activity of trypsin appears weaker than its canonical endopeptidase activity and exhibits a certain degree of sequence-dependency. This may explain the limited success and limited reproducibility across different proteomic studies aimed at identifying arginylated proteins. It is important to note that, while in our experimental conditions relatively high trypsin-to-substrate ratios were used to demonstrate a complete removal of N-terminal arginine in simplified systems, this requirement likely represents a conservative estimate relative to physiological proteomic samples. In complex biological samples, proteins bearing N-terminal arginine are expected to constitute a small fraction of the total protein. In conventional bottom-up proteomics workflows, which typically involve extended digestion periods, as internal lysine and arginine residues are progressively cleaved, it will result in a relative excess of active trypsin over the N-terminal arginine at the later stage. Furthermore, even in our simplified test conditions, noticeable removal of N-terminal arginine was observed at trypsin ratio as low as ∼1:500, which falls below typical trypsin concentrations in standard proteomic workflows. Therefore, the exopeptidase-like activity of trypsin is functional significant for applications requiring the preservation of N-terminal arginine.

N-terminal arginine, as products of Ate/AteL activity or proteolytic cleavage, is known to play important physiological roles in mediating protein turnover and in modulating protein functions. In numerous studies seeking to identify proteins with N-terminal arginine, many have relied on trypsin-based proteomics workflow, yet resulted in limited and/or poorly reproducible findings. Our results indicate that the underappreciated exopeptidase-like activity of trypsin may have contributed to these difficulties by erasing the N-terminal arginine. While the exopeptidase-like activity of trypsin appears to be relatively weak compared to its canonical endopeptidase activity, which likely allows the recovery of some N-terminal arginine residues in past studies, this activity would still substantially compromise sensitivity and robustness for this assay, potentially leading to a systematic underrepresentation of candidate substrates. Thus, the widespread use of trypsin in high throughput analyses may have obscured the prevalence of proteins or protein fragments bearing N-terminal arginine, leading to potential misinterpretations of the impacts of the corresponding protein modifications and/or proteases.

Our results further suggest that alternative digestion strategies should be considered when the goal is to identify proteins bearing N-terminal arginine, whether generated by arginylation or proteolytic cleavage. While Arg-C exhibits sequence-dependent, exopeptidase-like behavior, both Lys-C and LysargiNase preserve N-terminal arginine, making them suitable alternatives to trypsin or Arg-C. However, these proteases may introduce other technical trade-offs. For example, digestion with Lys-C alone typically produces longer peptides, which may complicate MS analysis. LysargiNase, by generating peptides with N-terminal arginine/lysine may enhance b-ion coverage, albeit often at the expense of y-ion detection. In addition, for native proteins whose N-termini lack a positively charged residue or modification, LysargiNase digestion would generate N-terminal peptides without a basic residue at either terminus, potentially resulting in lower ionization efficiency of these peptides. Depending on instrument sensitivity and the exact purpose, these concessions may be acceptable to retain N-terminal arginine. Additional enzymes, such as Glu-C or chymotrypsin, may also be considered, as they cleave C-terminal to residues other than arginine.

More broadly, our findings highlight the importance of carefully selecting proteases when studying protein N-termini. These include the analysis of N-terminal lysine, which can be generated by the action of proteases and may be equally important to N-terminal arginine as part of a type I N-degron or by affecting protein function. While this question is beyond the scope of the present investigation, it is conceivable that this residue can be also negatively affected by the usages of trypsin and even Lys-C. Therefore, interested investigators may need to consider confirming the suitability of the conventional digestion strategies and alternatives.

Taken together, our findings reveal a previously unrecognized limitation of trypsin in standard proteomics workflows and suggest that the widespread reliance on this enzyme may systematically compromise the detection of proteins bearing N-terminal basic residues. Re-evaluating digestion strategies may therefore be critical for accurately characterizing N-terminal modifications and proteolytic products in proteome-wide studies.

## Methods

### N-terminal arginine-specific antibodies

N-terminal arginine-specific antibodies were generated by GenScript^18,37,72^. Antibodies were raised in rabbits against peptides containing N-terminal arginine and cross adsorbed with peptides containing an identical primary sequence but without an N-terminal arginine.

Peptide antigen sequences:

(R)AA15: (R)AAVAAENEEIGAHIKHC
(R)-HIF: (R)EGAGGENEKKKMSC
(R)DD11: (R)DDIAALVVDC
(R)EHQ: (R)EHQLLC

Specificity of arginylation-specific antibodies were previously determined by ELISA and dot blotting for RAA15, R-HIF, RDD11, and REHQ^18,37,72^.

*Peptides*: all peptides were ordered from GenScript, with purities confirmed by MS to be greater than 90%.

### Protein purification

UBI_RDD-GFP-6xHis recombinant protein was expressed with IPTG induction in BL21 *E. coli* Codon plus strain in the presence of 100μg/mL ampicillin. Bacteria also constitutively co-expressed UBP1 to cleave the N-terminal ubiquitin to expose an N-terminal arginine as described by Wojtowicz, et al^73,74^. Recombinant protein was purified using native Ni-NTA purification and quantified by Bradford analysis. The purity of the protein and the presence of the N-terminal arginine were confirmed by Western blot and Coomassie stained SDS-PAGE, respectively.

### Digestion of peptides with trypsin or alternative peptidases

Mass spectrometry grade trypsin (ThermoFisher #90057) was reconstituted in 0.9mM EDTA in HBSS to 1.18ug/uL with a final working concentration of 1ug/uL. Cell-culture grade trypsin (0.25%,ThermoFisher #R001100) was used as-is or diluted in 0.9mM EDTA in HBSS before mixing with the peptide or protein solution, leading to a final concentration starting at ∼2.1ug/uL. N-terminally arginylated peptides, or their non-arginylated counterparts, were initially dissolved by DMSO into a concentration of 500ng/uL. For digestion, the peptides were added to the digestion buffer to a final working concentration of 75ng/uL in the reaction buffer (0.9mM EDTA in HBSS) and incubated at 37°C for the specified time with the respective trypsin treatment, or buffer alone when no trypsin was used. To inactivate trypsin, samples were heated to 100°C in a thermocycler with heated lid for 30 minutes. Recombinant Arg-C (PeproTech #450-54) was reconstituted to 1ug/uL in water. ArgC was previously lyophilized in 50mM Tris-HCl, pH 8.0 with 10mM CaCl_2_. Lys-C (New England Biolabs #P8109) was reconstituted to 100ng/uL in water. Lys-C was previously lyophilized in 100mM Tris-HCl, pH 8.0. LysargiNase (Sigma-Aldrich EMS0008) was reconstituted to 100ng/uL in water. LysargiNase was previously lyophilized in 500mM HEPES, 50mM CaCl_2,_ pH 7.5. Peptides were incubated with Arg-C, Lys-C, or LysargiNase at 37°C for the specified time in the matching buffer.

### Inhibition of trypsin

Cell culture-grade trypsin was mixed with trypsin neutralizing solution (TNS, Lonza Bioscience #CC-5002) at a ratio of 1:1 or 1: (trypsin to TNS; the manufacturer’s recommended ratio was 1:2) and incubated at room temperature for 15 minutes. 17uL of inhibited trypsin mixture was added to 3uL of 0.5mg/mL peptide substrates and incubated at 37°C for 30 minutes followed by heat-inactivation for 30 minutes at 100°C.

### Digestion of proteins with trypsin

The purified proteins (RDD-GFP-6xHis or Casein) were incubated with trypsin, with conditions similar as described with the peptide experiments.

### Dot blotting

Samples incubated with or without trypsin were pipetted in triplicate to nitrocellulose membranes. Dots were left to air dry for 90 minutes before washing twice with phosphate buffered saline (PBS). Membranes were briefly rinsed with PBST before shaking with Western Blocking Reagent (Roche #11921673001) for 1 hour at room temperature. Primary (arginine-specific) antibodies were diluted in blocking buffer and incubated with membranes overnight at 4°C. Membranes were washed with PBST, then incubated with goat-derived anti-rabbit-HRP (ThermoFisher #31460) for 1 hour at room temperature. Membranes were washed with PBST, then incubated with ECL substrate (ThermoFisher #32106) and imaged with an Amersham Imager 600 (GE Life Sciences).

### Mass spectrometry

Matrix assisted desorption/ionization time-of-flight (MALDI-TOF) analysis was conducted on a Bruker autoflex maX with reflectron mode set to 700-3500 Da. Samples were cleaned up with a ZipTip and eluted in 70:30(v/v) ACN:H2O +0.1% FA. Samples were mixed with a matrix of saturated CHCA (α-Cyano-4-hydroxycinnamic acid) prepared in 70:30(v/v) CAN:H2O +0.1% FA in a 1:1 ratio. The sample:matrix mixture was spotted on the MALDI plate (1 or 2µL) for analysis. For liquid chromatography coupled tandem mass spectrometry (LC-MS/MS), peptide samples were desalted using C18 SPE columns (Waters), completely dried in a speed vacuum centrifuge, and resuspended in 25ul 0.1% formic acid in water. Peptides were analyzed using a Q-Exactive Plus hybrid quadrupole Orbitrap (ThermoFisher) with a Vanquish Neo UHPLC (ThermoFisher). Peptides were separated on a 50cm C18 EZ-Spray column (ThermoFisher). Data-dependent acquisition was employed, selecting the Top20 most abundant peptide ions for MS/MS. Full MS Scans were acquired at 70,000 resolution. Raw data files were processed in MaxQuant (www.maxquant.org) against the provided peptides sequences supplemented with reverse and contaminant protein sequences.

### SDS PAGE

5uL of sample was diluted with an equal volume of 2X Laemmli loading buffer with 10% beta-mercaptoethanol, heated to 95°C for 10 minutes, and SDS PAGE was performed. Gels were stained with Colloidal Blue Staining kit (Invitrogen #LC6025) following manufacturer instructions.

## Acknowledgements

This work was supported by (NIGMS 1R01GM138557 and Sylvester Comprehensive Cancer Center Bridge Funding BFA-2025-04). MALDI-TOF analyses were performed at the University of Florida Mass Spectrometry Research and Education Center, which is in-part supported by NIH S10 OD0217558-01A1 and NIH S10 OD030250-01A1. We thank Dr. Dale Chaput and BioMS Discovery at the University of South Florida Advanced Research Core for Mass Spectrometry (ARC-MS) for performing LC-MS/MS.

## Abbreviations

Ate: arginyltransferase
AteL: Ate-like protein
CC: cell culture
GFP: green-fluorescence protein
LC-MS/MS: liquid chromatography coupled tandem mass spectrometry
MALDI-TOF: matrix-assisted laser desorption/ionization time-of-flight
MS: Mass Spectrometry
MS/MS: tandem MS
PAGE: Polyacrylamide gel electrophoresis

## Cited References

1. Baldwin, M. A. Protein Identification by Mass Spectrometry: Issues to be Considered *. Mol. Cell. Proteomics 3, 1–9 (2004).

2. Noor, Z., Ahn, S. B., Baker, M. S., Ranganathan, S. & Mohamedali, A. Mass spectrometry–based protein identification in proteomics—a review. Brief. Bioinform. 22, 1620–1638 (2021).

3. Vajda, T. & Szabó, T. Specificity of trypsin and alpha-chymotrypsin towards neutral substrates. Acta Biochim. Biophys. Acad. Sci. Hung. 11, 287–294 (1976).

4. Olsen, J. V., Ong, S.-E. & Mann, M. Trypsin Cleaves Exclusively C-terminal to Arginine and Lysine Residues*. Mol. Cell. Proteomics 3, 608–614 (2004).

5. Burkhart, J. M., Schumbrutzki, C., Wortelkamp, S., Sickmann, A. & Zahedi, R. P. Systematic and quantitative comparison of digest efficiency and specificity reveals the impact of trypsin quality on MS-based proteomics. J. Proteomics 75, 1454–1462 (2012).

6. Ferber, S. & Ciechanover, A. Role of arginine-tRNA in protein degradation by the ubiquitin pathway. Nature 326, 808–811 (1987).

7. Varshavsky, A. The N-end rule: functions, mysteries, uses. Proc. Natl. Acad. Sci. 93, 12142–12149 (1996).

8. Tasaki, T., et al. The substrate recognition domains of the N-end rule pathway. J. Biol. Chem. 284, 1884–1895 (2009).

9. Varshavsky, A. The N-end rule pathway and regulation by proteolysis. Protein Sci. Publ. Protein Soc. 20, 1298–1345 (2011).

10. Karakozova, M., et al. Arginylation of β-Actin Regulates Actin Cytoskeleton and Cell Motility. Science 313, 192–196 (2006).

11. Rai, R., et al. Arginyltransferase regulates alpha cardiac actin function, myofibril formation and contractility during heart development. Dev. Camb. Engl. 135, 3881–3889 (2008).

12. Zhang, F., Saha, S. & Kashina, A. Arginylation-dependent regulation of a proteolytic product of talin is essential for cell–cell adhesion. J. Cell Biol. 197, 819–836 (2012).

13. Momose, K. & Kaji, A. Soluble amino acid-incorporating system. 3. Further studies on the product and its relation to the ribosomal system for incorporation. J. Biol. Chem. 241, 3294–3307 (1966).

14. Soffer, R. L. The arginine transfer reaction. Biochim. Biophys. Acta BBA - Nucleic Acids Protein Synth. 155, 228–240 (1968).

15. Balzi, E., Choder, M., Chen, W. N., Varshavsky, A. & Goffeau, A. Cloning and functional analysis of the arginyl-tRNA-protein transferase gene ATE1 of Saccharomyces cerevisiae. J. Biol. Chem. 265, 7464–7471 (1990).

16. Kwon, Y. T., Kashina, A. S. & Varshavsky, A. Alternative Splicing Results in Differential Expression, Activity, and Localization of the Two Forms of Arginyl-tRNA-Protein Transferase, a Component of the N-End Rule Pathway. Mol. Cell. Biol. 19, 182–193 (1999).

17. Hu, R.-G. et al. Arginyltransferase, Its Specificity, Putative Substrates, Bidirectional Promoter, and Splicing-derived Isoforms *. J. Biol. Chem. 281, 32559–32573 (2006).

18. Moorthy, B. T., et al. The evolutionarily conserved Arginyltransferase1 mediates a pVHL-independent oxygen-sensing pathway in mammalian cells. Dev. Cell 57, 654–669.e9 (2022).

19. Tompa, P., et al. On the Sequential Determinants of Calpain Cleavage*. J. Biol. Chem. 279, 20775–20785 (2004).

20. Piatkov, K. I., Oh, J.-H., Liu, Y. & Varshavsky, A. Calpain-generated natural protein fragments as short-lived substrates of the N-end rule pathway. Proc. Natl. Acad. Sci. 111, E817–E826 (2014).

21. Shinkai-Ouchi, F., et al. Predictions of Cleavability of Calpain Proteolysis by Quantitative Structure-Activity Relationship Analysis Using Newly Determined Cleavage Sites and Catalytic Efficiencies of an Oligopeptide Array *. Mol. Cell. Proteomics 15, 1262–1280 (2016).

22. Stennicke, H. R., Renatus, M., Meldal, M. & Salvesen, G. S. Internally quenched fluorescent peptide substrates disclose the subsite preferences of human caspases 1, 3, 6, 7 and 8. Biochem. J. 350, 563–568 (2000).

23. Timmer, J. C. & Salvesen, G. S. Caspase substrates. Cell Death Differ. 14, 66–72 (2007).

24. Shen, J., et al. Caspase-1 recognizes extended cleavage sites in its natural substrates. Atherosclerosis 210, 422–429 (2010).

25. Rai, R., et al. Arginyltransferase ATE1 suppresses cell tumorigenic potential and inversely correlates with metastases in human cancers. Oncogene 35, 4058–4068 (2016).

26. Birnbaum, M. D., et al. Reduced Arginyltransferase 1 is a driver and a potential prognostic indicator of prostate cancer metastasis. Oncogene 38, 838–851 (2019).

27. Xu, C., Li, Y.-M., Sun, B., Zhong, F.-J. & Yang, L.-Y. ATE1 Inhibits Liver Cancer Progression through RGS5-Mediated Suppression of Wnt/β-Catenin Signaling. Mol. Cancer Res. 19, 1441–1453 (2021).

28. Lazar, I., et al. Arginyl-tRNA-protein transferase 1 (ATE1) promotes melanoma cell growth and migration. FEBS Lett. 596, 1468–1480 (2022).

29. Nawale, L., et al. ATE1 promotes breast cancer progression via arginylation-dependent regulation of MAPK-MYC signaling. Cell Commun. Signal. 23, 390 (2025).

30. Kwon, Y. T., et al. An essential role of N-terminal arginylation in cardiovascular development. Science 297, 96–99 (2002).

31. Kurosaka, S., et al. Arginylation regulates myofibrils to maintain heart function and prevent dilated cardiomyopathy. J. Mol. Cell. Cardiol. 53, 333–341 (2012).

32. Lian, L., et al. Loss of ATE1-mediated arginylation leads to impaired platelet myosin phosphorylation, clot retraction, and in vivo thrombosis formation. Haematologica 99, 554–560 (2014).

33. Singh, K., et al. Arginyltransferase knockdown attenuates cardiac hypertrophy and fibrosis through TAK1-JNK1/2 pathway. Sci. Rep. 10, 598 (2020).

34. Masdehors, P., Glaisner, S., Maciorowski, Z., Magdelénat, H. & Delic, J. Ubiquitin-dependent protein processing controls radiation-induced apoptosis through the N-end rule pathway. Exp. Cell Res. 257, 48–57 (2000).

35. Decca, M. B., et al. Post-translational arginylation of calreticulin: a new isospecies of calreticulin component of stress granules. J. Biol. Chem. 282, 8237–8245 (2007).

36. López Sambrooks, C., Carpio, M. A. & Hallak, M. E. Arginylated calreticulin at plasma membrane increases susceptibility of cells to apoptosis. J. Biol. Chem. 287, 22043–22054 (2012).

37. Kumar, A., et al. Posttranslational arginylation enzyme Ate1 affects DNA mutagenesis by regulating stress response. Cell Death Dis. 7, e2378 (2016).

38. Deka, K., Singh, A., Chakraborty, S., Mukhopadhyay, R. & Saha, S. Protein arginylation regulates cellular stress response by stabilizing HSP70 and HSP40 transcripts. Cell Death Discov. 2, 16074 (2016).

39. Zanakis, M. F., Chakraborty, G., Sturman, J. A. & Ingoglia, N. A. Posttranslational Protein Modification by Amino Acid Addition in Intact and Regenerating Axons of the Rat Sciatic Nerve. J. Neurochem. 43, 1286–1294 (1984).

40. Shyne-Athwal, S., Riccio, R. V., Chakraborty, G. & Ingoglia, N. A. Protein Modification by Amino Acid Addition Is Increased in Crushed Sciatic But Not Optic Nerves. Science 231, 603–605 (1986).

41. Chakraborty, G. & Ingoglia, N. A. N-terminal arginylation and ubiquitin-mediated proteolysis in nerve regeneration. Brain Res. Bull. 30, 439–445 (1993).

42. Wang, Y. M. & Ingoglia, N. A. N-terminal arginylation of sciatic nerve and brain proteins following injury. Neurochem. Res. 22, 1453–1459 (1997).

43. Brower, C. S. & Varshavsky, A. Ablation of arginylation in the mouse N-end rule pathway: loss of fat, higher metabolic rate, damaged spermatogenesis, and neurological perturbations. PloS One 4, e7757 (2009).

44. Galiano, M. R., Goitea, V. E. & Hallak, M. E. Post-translational protein arginylation in the normal nervous system and in neurodegeneration. J. Neurochem. 138, 506–517 (2016).

45. Kim, E., Kim, S., Lee, J. H., Kwon, Y. T. & Lee, M. J. Ablation of Arg-tRNA-protein transferases results in defective neural tube development. BMB Rep. 49, 443–448 (2016).

46. Wang, J., et al. Arginyltransferase ATE1 is targeted to the neuronal growth cones and regulates neurite outgrowth during brain development. Dev. Biol. 430, 41–51 (2017).

47. Xu, Z., Payoe, R. & Fahlman, R. P. The C-terminal Proteolytic Fragment of the Breast Cancer Susceptibility Type 1 Protein (BRCA1) Is Degraded by the N-end Rule Pathway*. J. Biol. Chem. 287, 7495–7502 (2012).

48. Piatkov, K. I., Brower, C. S. & Varshavsky, A. The N-end rule pathway counteracts cell death by destroying proapoptotic protein fragments. Proc. Natl. Acad. Sci. U. S. A. 109, E1839–E1847 (2012).

49. Brower, C. S., Piatkov, K. I. & Varshavsky, A. Neurodegeneration-Associated Protein Fragments As Short-Lived Substrates of the N-End Rule Pathway. Mol. Cell 50, 161–171 (2013).

50. Wadas, B., Piatkov, K. I., Brower, C. S. & Varshavsky, A. Analyzing N-terminal Arginylation through the Use of Peptide Arrays and Degradation Assays. J. Biol. Chem. 291, 20976–20992 (2016).

51. Wang, J., et al. Target site specificity and in vivo complexity of the mammalian arginylome. Sci. Rep. 8, 16177 (2018).

52. Decca, M. B., et al. Protein Arginylation in Rat Brain Cytosol: A Proteomic Analysis. Neurochem. Res. 31, 401–409 (2006).

53. Wong, C. C. L., et al. Global Analysis of Posttranslational Protein Arginylation. PLOS Biol. 5, e258 (2007).

54. Lin, Z., et al. An unbiased proteomic platform for ATE1-based arginylation profiling. Nat. Chem. Biol. 21, 1970–1980 (2025).

55. Davydov, I. V. & Varshavsky, A. RGS4 Is Arginylated and Degraded by the N-end Rule Pathway in Vitro *. J. Biol. Chem. 275, 22931–22941 (2000).

56. Lee, M. J., et al. RGS4 and RGS5 are in vivo substrates of the N-end rule pathway. Proc. Natl. Acad. Sci. U. S. A. 102, 15030–15035 (2005).

57. Seo, T., et al. R-catcher, a potent molecular tool to unveil the arginylome. Cell. Mol. Life Sci. CMLS 78, 3725–3741 (2021).

58. Gevaert, K., et al. Exploring proteomes and analyzing protein processing by mass spectrometric identification of sorted N-terminal peptides. Nat. Biotechnol. 21, 566–569 (2003).

59. Arnesen, T., et al. Proteomics analyses reveal the evolutionary conservation and divergence of N-terminal acetyltransferases from yeast and humans. Proc. Natl. Acad. Sci. 106, 8157–8162 (2009).

60. Helbig, A. O., et al. Profiling of N-Acetylated Protein Termini Provides In-depth Insights into the N-terminal Nature of the Proteome *. Mol. Cell. Proteomics 9, 928–939 (2010).

61. Mommen, G. P. M., et al. Unbiased Selective Isolation of Protein N-terminal Peptides from Complex Proteome Samples Using Phospho Tagging (PTAG) and TiO2-based Depletion. Mol. Cell. Proteomics 11, 832–842 (2012).

62. Frey, A. M., Chaput, D. & Shaw, L. N. Insight into the human pathodegradome of the V8 protease from Staphylococcus aureus. Cell Rep. 35, 108930 (2021).

63. Britton, M., et al. Selective inhibitor of proteasome’s caspase-like sites sensitizes cells to specific inhibition of chymotrypsin-like sites. Chem. Biol. 16, 1278–1289 (2009).

64. Kraus, M., et al. The novel β2-selective proteasome inhibitor LU-102 synergizes with bortezomib and carfilzomib to overcome proteasome inhibitor resistance of myeloma cells. Haematologica 100, 1350–1360 (2015).

65. Ju, S., Cha-Molstad, H. & Lee, C. Tracing the mark of arginine. Nat. Chem. Biol. 21, 1837–1838 (2025).

66. Norrgran, J., et al. Optimization of digestion parameters for protein quantification. Anal. Biochem. 393, 48–55 (2009).

67. Getie-Kebtie, M., Sultana, I., Eichelberger, M. & Alterman, M. Label-free mass spectrometry-based quantification of hemagglutinin and neuraminidase in influenza virus preparations and vaccines. Influenza Other Respir. Viruses 7, 521–530 (2013).

68. Egeland, S. V., Reubsaet, L. & Halvorsen, T. G. The pros and cons of increased trypsin-to-protein ratio in targeted protein analysis. J. Pharm. Biomed. Anal. 123, 155–161 (2016).

69. Optimal conditions for carrying out trypsin digestions on complex proteomes: From bulk samples to single cells. J. Proteomics 297, 105109 (2024).

70. Swaney, D. L., Wenger, C. D. & Coon, J. J. Value of Using Multiple Proteases for Large-Scale Mass Spectrometry-Based Proteomics. J. Proteome Res. 9, 1323–1329 (2010).

71. Wu, Z., Huang, J., Huang, J., Li, Q. & Zhang, X. Lys-C/Arg-C, a More Specific and Efficient Digestion Approach for Proteomics Studies. Anal. Chem. 90, 9700–9707 (2018).

72. Zhang, F., Saha, S. & Kashina, A. Arginylation-dependent regulation of a proteolytic product of talin is essential for cell–cell adhesion. J. Cell Biol. 197, 819–836 (2012).

73. Wojtowicz, A., et al. Expression of yeast deubiquitination enzyme UBP1 analogues in E. coli. Microb. Cell Factories 4, 17 (2005).

74. Wojtowicz-Krawiec, A., et al. Use of Ubp1 protease analog to produce recombinant human growth hormone in Escherichia coli. Microb. Cell Factories 13, 113 (2014).

